# A giant leap in sequence space reveals the intracellular complexities of evolving a new function

**DOI:** 10.1101/2020.05.27.118489

**Authors:** Kelsi R. Hall, Katherine J. Robins, Michelle H. Rich, Mark J. Calcott, Janine N. Copp, Elsie M. Williams, Rory F. Little, Ralf Schwörer, Gary B. Evans, Wayne M. Patrick, David F. Ackerley

## Abstract

Selection for a promiscuous enzyme activity provides substantial opportunity for competition between endogenous and new substrates to influence the evolutionary trajectory, an aspect that has generally been overlooked in laboratory directed evolution studies. We evolved the *Escherichia coli* nitro/quinone reductase NfsA to detoxify chloramphenicol by randomising eight active site residues simultaneously and interrogating ∼250,000,000 reconfigured NfsA variants. Analysis of every possible evolutionary intermediate of the two best chloramphenicol reductases revealed complex epistatic interactions that restrict each hypothetical trajectory. In both cases, improved chloramphenicol detoxification was only possible after one essential substitution had eliminated activity with endogenous quinone substrates. Unlike the predominantly weak trade-offs seen in previous experimental studies, this substrate incompatibility suggests endogenous metabolites have considerable potential to shape evolutionary outcomes. Unselected prodrug-converting activities were mostly unaffected, which emphasises the importance of negative selection to effect enzyme specialisation, and offers an application for the evolved genes as dual-purpose selectable/counter-selectable markers.

## Introduction

Many (if not all) enzymes are promiscuous, meaning that in addition to their primary biological role(s) they can catalyse minor side reactions that have no apparent physiological relevance, either because they are too inefficient or because the substrate is not naturally encountered (Copley, 2015). From an evolutionary perspective, promiscuity can play an important role in contingency, i.e. providing a reservoir of potential functions that a cell can tap in response to changing circumstances (O’Brien and Herschlag, 1999; Copley, 2015). As demonstrated by the emergence of resistance to xenobiotic pollutants or clinical antibiotics (O’Brien and Herschlag, 1999; Hall, 2004; Ramos *et al.*, 2005; Copley, 2009; Khersonsky and Tawfik, 2010), a strong selection pressure can cause latent promiscuous activities to be rapidly amplified to physiologically relevant levels (Newton *et al.*, 2015).

Catalytic transitions to an alternate substrate have been modelled experimentally using iterative rounds of random mutagenesis (e.g., error-prone PCR), a powerful directed evolution strategy that enables adaptive landscapes to be explored under defined laboratory conditions (Kaltenbach *et al.*, 2015; Kaltenbach *et al.*, 2016). These laboratory evolution studies have indicated that selection for substantial increases in a promiscuous activity typically results in only weak trade-offs against the native activity; and therefore, the transition from one primary function to another tends to progress via generalist enzyme intermediates (Kaltenbach *et al.*, 2016). Two leading teams have offered contrasting hypotheses to explain this phenomenon. In 2005, Tawfik and co-workers proposed that enzymes possess an innate ‘robustness’ and stability that buffers them against the potentially detrimental effects of novel mutations, coupled with a ‘plasticity’ that can amplify promiscuous functions with relatively few mutations (Aharoni *et al.*, 2005). More recently, Tokuriki and co-workers demonstrated that a robust native activity is not a prerequisite for weak trade-offs, and suggested that their predominance in the literature may instead be artefactual; a consequence of laboratory evolution studies being highly biased toward strong selection for a new function without any selection against the native activity (Kaltenbach *et al.*, 2016). They argue that it is unclear how specialisation can occur in this manner, and that in nature, selection might frequently exist to erode the original function. By way of example, they offer a scenario where the native and new substrate compete for the same active site (Kaltenbach *et al.*, 2016).

In addition to their exclusive emphasis on positive selection, we note that these pioneering previous studies were also heavily biased toward heterologous enzyme expression and/or a transition in activity from one exogenously applied substrate to another (Kaltenbach *et al.*, 2016). Thus, there has been little consideration of how the native substrate might influence the evolutionary trajectory. We were therefore motivated to study evolution of a promiscuous function within the native host environment, and were particularly interested to focus on the key catalytic changes driving the transition. Recognising that the stochastic nature of iterative random mutagenesis is unlikely to yield the most efficient pathway to an evolved outcome, we sought to implement simultaneous mass-mutagenesis on a massive scale that would allow us to retrospectively assess all possible intermediates of our top variants, and infer the most plausible evolutionary trajectories. We were able to achieve both these goals by employing the *Escherichia coli* nitro/quinone reductase NfsA as a new model system that offers several key advantages. NfsA is a member of a large bacterial superfamily comprising highly promiscuous FMN-dependent oxidoreductases that accept electrons from NAD(P)H and transfer them to a diverse range of substrates (Williams *et al.*, 2015; Akiva *et al.*, 2017). Expression of *nfsA* is governed by the *soxRS* regulon, and NfsA is thought to guard against oxidative stress through reduction of water-soluble quinones such as 1,4-benzoquinone (Liochev *et al.*, 1999; Paterson *et al.*, 2002; Copp *et al.*, 2017). Although most efficient with quinone substrates, NfsA is also able to reduce a wide diversity of nitroaromatic compounds (Valiauga *et al.*, 2017). This is generally believed to represent non-physiological substrate ambiguity, as there are relatively few nitroaromatic natural products, and in many cases nitro-reduction yields a more toxic derivative (Winkler and Hertweck, 2007; Parry *et al.*, 2011; Williams *et al.*, 2015). An important exception is that nitro-reduction of chloramphenicol transforms this antibiotic to a product that is not discernibly toxic to bacteria (Yunis, 1988; Smith *et al.*, 2007; Crofts *et al.*, 2019). We have observed that over-expressed native NfsA confers only slight chloramphenicol protection to *E. coli* host cells, but reasoned that we could select for improved detoxification in an extremely high throughput manner by plating variant libraries on chloramphenicol-amended media. Because members of the bacterial nitroreductase superfamily appear to have unusually plastic active sites (Akiva *et al.*, 2017), we considered that simultaneous mass-mutagenesis of up to eight active site residues should be possible. In effect, we aimed to strip NfsA of its engine, and then select for a superior configuration of parts assembled within the empty chassis. By leaping directly to a new fitness peak, we considered that we might arrive at synergistic combinations of substitutions that would be difficult to achieve by iterative random mutagenesis approaches.

## Results

### Design of an eight randomised codon nfsA gene library

We have previously conducted several different mutagenesis studies on *nfsA*, seeking to enhance activity with prodrugs and/or positron emission tomography (PET) imaging probes for cancer gene therapy applications (Williams, 2013; Copp *et al.*, 2017; Rich, 2017), or to assess potential collateral sensitivities between niclosamide and the antibiotics metronidazole and nitrofurantoin (Copp *et al.*, 2020). Based on this previous work we empirically identified eight active site residues (S41, L43, H215, T219, K222, S224, R225 and F227; Fig. 1A) as being individually mutable and having the potential to contribute to generically improved nitroreductase activity. We then designed a degenerate gene library to enable simultaneous randomisation of each residue. As complete randomisation of target codons (e.g., NNK degeneracy) would have yielded an impractically large library of >10^12^ (32^8^ or more) gene variants, we instead used a restricted set (Fig. 1B). The degenerate codon NDT was preferred at most sites, as this specifies 12 different amino acids that represent a balanced portfolio of small & large, polar & non-polar, aliphatic & aromatic, and negatively & positively charged side chains (Reetz *et al.*, 2008). However, at positions 219 and 222, NDT codons did not include the native NfsA residue as an option, so the alternative degeneracies NHT (12 unique amino acids) and VNG (11 unique amino acids) were chosen as acceptably balanced alternatives (Fig. 1B). In total, our library represented 430 million possible gene combinations, collectively specifying 394 million different NfsA variants.

**Figure 1:**
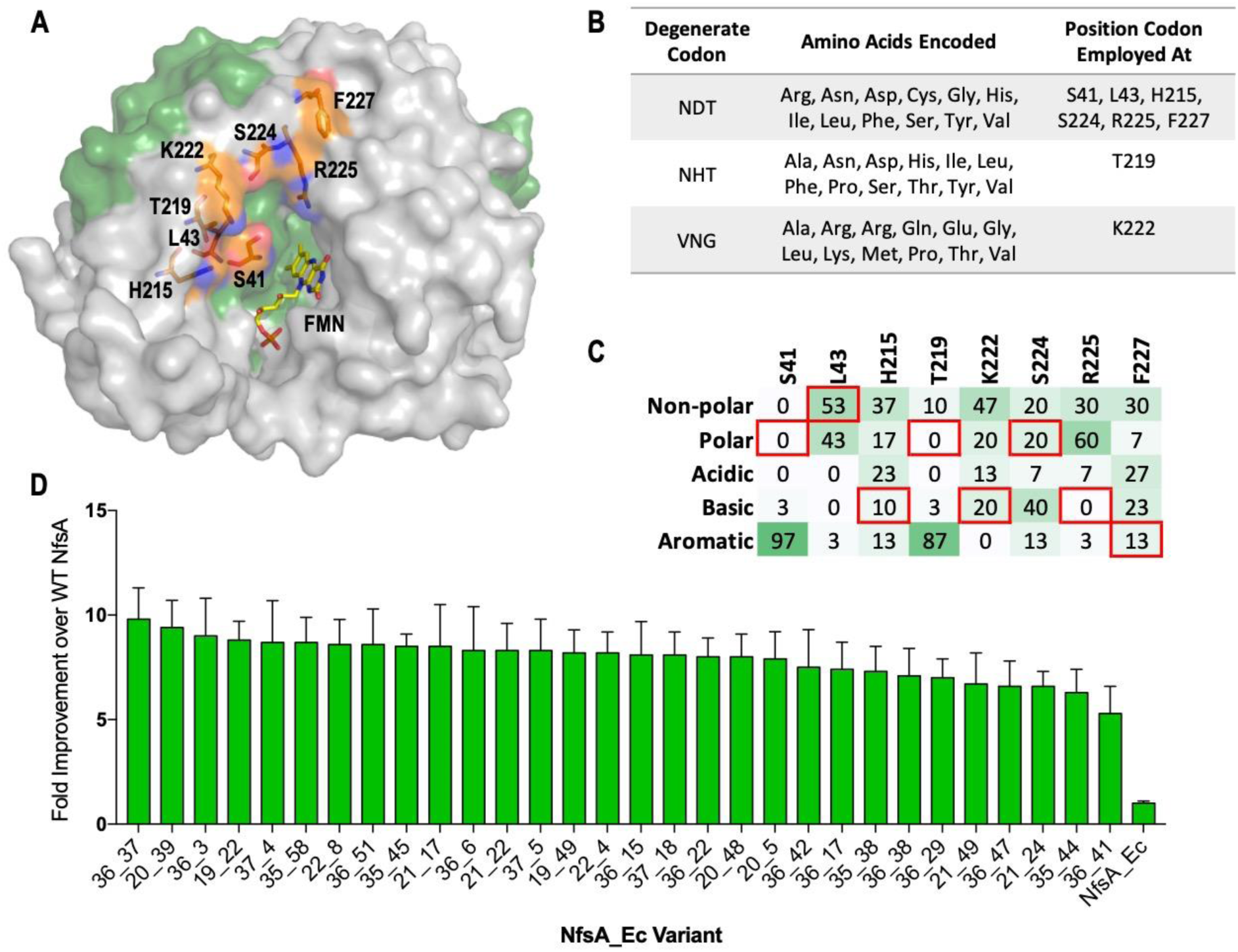
Creation, selection and characterisation of 30 top chloramphenicol detoxifying NfsA variants. (A) Structure of NfsA, based on PDB 1f5v. One monomer is shown is shown in grey and one monomer in green. The eight residues simultaneously targeted in NfsA (carbons in orange) and the FMN cofactor (carbons in yellow) are shown in stick form. For clarity, only one of the two FMN-binding active sites in the enzyme homodimer is portrayed. (B) Summary of the amino acid repertoire encoded by each degenerate codon. (C) Percentage of the five amino acid side-chain categories at each of the eight targeted positions for the top 30 chloramphenicol reducing variants. The property of the native amino acid at each position is boxed in red. (D) Fold-improvement in chloramphenicol EC_50_ values for *E. coli* 7NT strains expressing the top 30 chloramphenicol-detoxifying *nfsA* variants over the native *nfsA* control (far right). Data presented in Figures 1D represents the average of at least four biological repeats ± 1 S.D.

### Selection and characterisation of superior chloramphenicol-detoxifying NfsA variants

Following artificial synthesis and cloning, our library was used to transform *E. coli* 7NT cells (a strain in which endogenous nitroreductase genes had been deleted). We ultimately recovered a total of 398 million transformed colonies, a collection predicted by GLUE (Patrick *et al.*, 2003) to represent 252 million different NfsA variants. Despite the drastic reconfiguration of their encoded active sites, a surprising 0.05% of the gene variants (∼200,000 clones) were more effective than wild type *nfsA* (i.e., enabled colony formation on LB agar amended with 3 µM chloramphenicol, the lowest concentration at which wild type *nfsA* was unable to support host cell growth). This robust tolerance to active site randomisation confirmed that NfsA exhibits a substantial degree of active site plasticity.

We next plated the library on ≥45 µM chloramphenicol, recovering a total of 365 colonies. Retransformation of the variant-encoding plasmids into fresh 7NT host cells to eliminate any selected chromosomal mutations, followed by validation of activities in liquid growth assays, then sequencing and elimination of duplicates, yielded 30 top variants that exhibited evidence of a conserved genetic response to the chloramphenicol selection. Particularly strong trends were observed at positions 41 and 219 (where the native serine or threonine was substituted by an aromatic residue in ≥26 of the 30 cases), and at position 225 (100% substitution of the native arginine by an uncharged or acidic residue) (Fig. 1C). Only at position 43 was the native or functionally similar residue frequently retained (16/30 cases). In EC_50_ assays, 7NT cells expressing these 30 variants demonstrated nearly 6- to 10-fold greater chloramphenicol tolerance than those expressing native *nfsA* (Fig. 1D).

To evaluate the impact of the active site reconfiguration on catalytic activity, the top five chloramphenicol detoxifying NfsA variants were purified as His_6_-tagged proteins and evaluated in steady-state kinetics assays. We were surprised to discover that these variants exhibited only marginal (at most 2.2-fold) improvements in chloramphenicol *k*_*cat*_/*K*_*M*_ over wild type NfsA (Table 1). However, in every case the variants were impaired in *k*_*cat*_ (6-10 fold lower than NfsA) but greatly improved in *K*_*M*_ (8-13 fold lower than NfsA). Thus, it appeared that the *in vivo* improvements in chloramphenicol detoxification were driven primarily by enhanced substrate affinity.

**Table 1:**
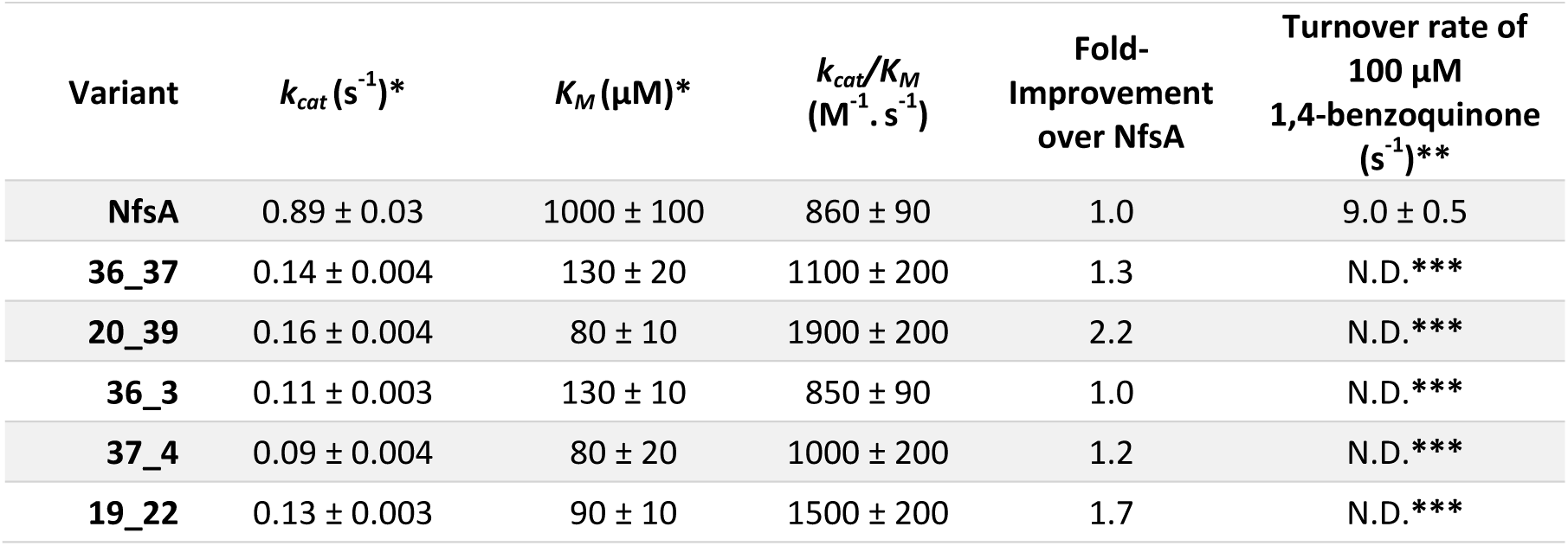
Kinetic parameters of chloramphenicol and 1,4*-*benzoquinone reduction for purified NfsA variants (the top five by *in vivo* EC_50_ ranking). Apparent *K*_*M*_ and *k*_*cat*_ at 250 µM NADPH were calculated using Graphpad 8.0. Kinetic parameters could not be accurately determined for 1,4*-*benzoquinone for any of the selected variants; in an attempt to detect trace activities, the catalytic rate of 1,4-benzoquinone reduction was measured at a single high concentration of 1,4-benzoquinone (100 µM), with reactions initiated by addition of 250 µM NADPH. All reactions were measured in triplicate and errors are ± 1 S.D. **\***Apparent *k*_*cat*_ and *K*_*M*_ as determined at 250 µM NADPH. **\*\***Measured rates following addition of 250 µM NADPH. **\*\*\***N.D. = not detectable (<0.1 s^-1^).

| Variant | $k_{cat}$ (s<sup>-1</sup>)* | $K_M$ (μM)* | $k_{cat}/K_M$<br>(M<sup>-1</sup> · s<sup>-1</sup>) | Fold-Improvement<br>over NfsA | Turnover rate of<br>100 μM<br>1,4-benzoquinone<br>(s<sup>-1</sup>)** |
| --- | --- | --- | --- | --- | --- |
| NfsA | 0.89 ± 0.03 | 1000 ± 100 | 860 ± 90 | 1.0 | 9.0 ± 0.5 |
| 36_37 | 0.14 ± 0.004 | 130 ± 20 | 1100 ± 200 | 1.3 | N.D.*** |
| 20_39 | 0.16 ± 0.004 | 80 ± 10 | 1900 ± 200 | 2.2 | N.D.*** |
| 36_3 | 0.11 ± 0.003 | 130 ± 10 | 850 ± 90 | 1.0 | N.D.*** |
| 37_4 | 0.09 ± 0.004 | 80 ± 20 | 1000 ± 200 | 1.2 | N.D.*** |
| 19_22 | 0.13 ± 0.003 | 90 ± 10 | 1500 ± 200 | 1.7 | N.D.*** |

We previously observed a similar phenomenon when using an error prone PCR strategy to evolve NfsA for improved activation of the anti-cancer prodrug PR-104A, with all top variants exhibiting a lower *k*_*cat*_ and lower *K*_*M*_ for PR-104A, and none being significantly improved in *k*_*cat*_/*K*_*M*_ over the native enzyme (Copp *et al.*, 2017). In that study, we postulated that the improved *in vivo* activities were a consequence of diminished competitive inhibition by native quinone substrates present in the *E. coli* cytoplasm; although the top variant was still active with 1,4-benzoquinone, we found its PR-104A reduction activity was less affected by addition of 1,4-benzoquinone to the reaction mix than was the case for wild type NfsA (Copp *et al.*, 2017). We were therefore interested to discover whether our improved chloramphenicol-detoxifying variants from the present study were impaired in 1,4- benzoquinone reduction. In all cases, we found that 1,4-benzoquinone reduction was unmeasurable (Table 1), suggesting that this activity had been actively and strongly selected against.

### Recreating all possible evolutionary trajectories for the top chloramphenicol detoxifying NfsA variants

We next sought to probe the contributions to improved chloramphenicol detoxification and/or diminished 1,4-benzoquinone reduction made by key substitutions, or combinations thereof. The top two chloramphenicol detoxifying variants (36_37 and 20_39) each had seven substitutions at the eight targeted positions, with both containing the wild-type residue leucine at position 43 (36_37 = S41Y, H215C, T219Y, K222V, S224R, R225V, F227G; 20_39 = S41Y, H215N, T219Y, K222R, S224Y, R225D, F227H; Figure 2A-C). Therefore, there are 126 possible intermediate forms (2^7^ – 2) between wild type NfsA and the evolved variant. Genes encoding the 126 intermediate forms for each variant were artificially synthesised, cloned and expressed in *E. coli* 7NT cells, and the chloramphenicol tolerance of the resulting strains assessed in EC_50_ growth assays. A Python script was generated to delineate all 5040 (7!) possible evolutionary trajectories and the output used to generate full network graphs (Figure 2D-E, Supplementary Figure S1). We then considered whether traditional stepwise directed evolution strategies, which require each substitution to directly improve the selected activity (e.g., across iterative rounds of error prone PCR), could have plausibly generated either of variants 36_37 or 20_39. For the purposes of this analysis we considered “improvement” to be a >16% increase in chloramphenicol EC_50_ for each step, as this was the average error across all EC_50_ measurements. In neither case was there a clear path from NfsA to the final variant that involved exclusively upward steps in the hypothetical evolutionary trajectory (Figure 2D-F). Nevertheless, it was evident that the final two substitutions (H215C and K222V for variant 36_37, and K222R and S224Y for variant 20_39) did not contribute substantially to the overall chloramphenicol detoxifying activity of each variant. Thus, we concluded that iterative evolutionary strategies could have plausibly generated NfsA variants exhibiting similar levels of chloramphenicol detoxifying activity to 36_37 and 20_39, but that there were very few accessible pathways for this (Figure 2D-E).

**Figure 2:**
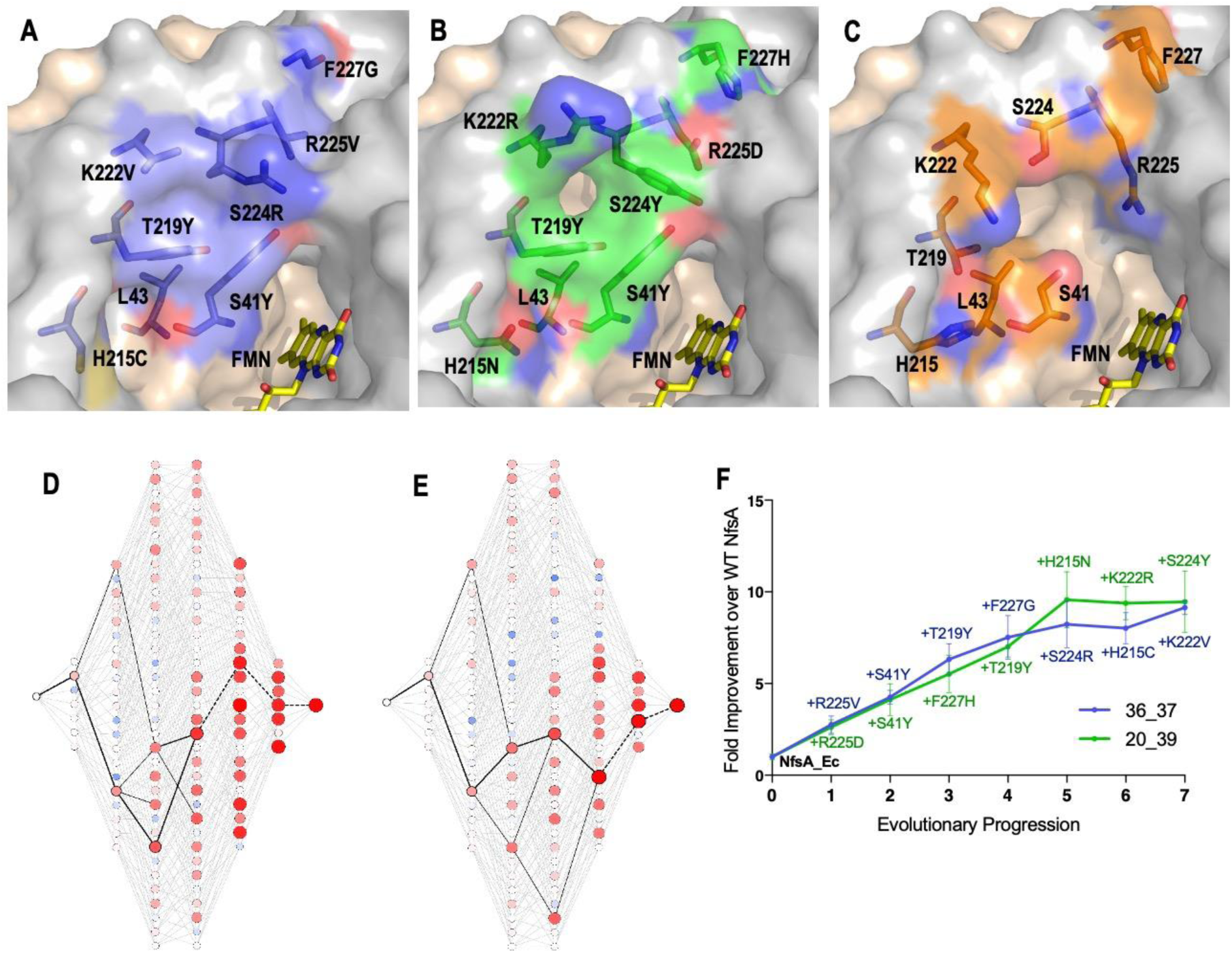
Recreating the hypothetical evolutionary trajectories of NfsA variants 36_37 and 20_39. (A-C) Residues in the active site of 36_37 (A, blue), 20_39 (B, green) and wild-type NfsA (C, orange); based on PDB 1f5v (Kobori *et al.*, 2001). In each panel, one NfsA monomer is shaded grey and the other is pale pink. The orientation of the mutated residues in 36_37 and 20_39 was predicted using the mutagenesis wizard on PyMOL, which selected the most likely rotamer conformation based on the frequencies of occurrence in proteins while avoiding clashes with other residues. (D-E) All 5040 possible evolutionary trajectories of 36_37 (D) and 20_39 (E). Black lines represent primary paths in which each step resulted in a ≥ 16% increase in chloramphenicol detoxification. Thick black lines represent the most probable stepwise evolutionary trajectory as explained in Figure F. The colour and diameter of nodes corresponds to the fold-improvement in chloramphenicol detoxification over wild-type NfsA (blue/smaller = less active, red/larger = more active). A larger version of each image is provided in Supplementary Figure S1. (F) The most plausible stepwise evolutionary trajectory for each of variant 36_37 (blue) and variant 20_39 (green). To establish these, the substitution which resulted in the greatest improvement in chloramphenicol detoxification was selected at each point in the evolutionary progression. If no substitutions improved chloramphenicol detoxification, then the substitution was selected which resulted in the smallest decrease in activity (shown as a dotted black light in D-E). Data presented in Figures 2D-F represent the average of at least four biological repeats ± 1 S.D.

The dearth of accessible evolutionary pathways suggested extensive epistasis, a phenomenon that several teams have previously observed when evolving enzymes (Weinreich *et al.*, 2005; Poelwijk *et al.*, 2011; Kaltenbach and Tokuriki, 2014; Yang *et al.*, 2019; Ben-David *et al.*, 2020), where the fitness effects of certain substitutions only manifest when other substitutions have already been made. Most prominently, we noted that only one of the seven substitutions present in each of variants 36_37 and 20_39 significantly enhanced chloramphenicol detoxification when introduced on an individual basis (Figure 3A).

**Figure 3:**
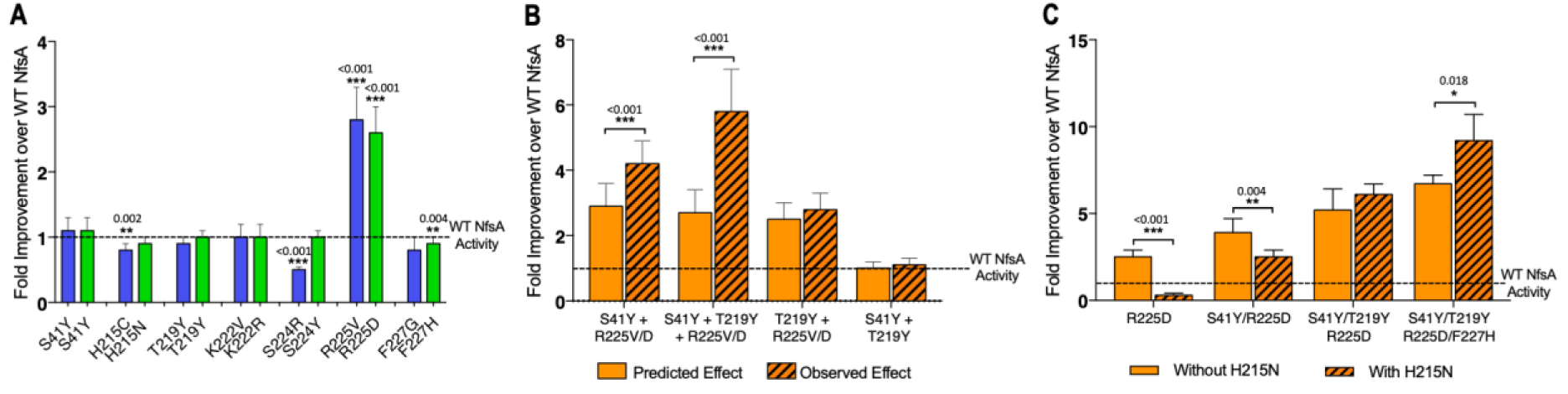
Complex epistatic interactions exist in 36-37 and 20_39. (A) The effect on chloramphenicol detoxification of introducing individual substitutions present in 36_37 (blue) and 20_39 (green) into wild type (WT) NfsA. (B) Observation of epistatic interactions between S41Y, T219Y and R225V/D. The predicted multiplicative effects (solid bars) were calculated by multiplying the fold-increase conferred by individual amino acid substitutions. The error of the predicted effects were derived using an error propagation equation 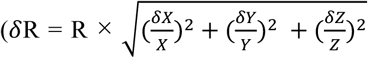 where *δ*X, *δ*Y, *δ*Z is the error of EC_50_ values X, Y and Z and *δ*R is the calculated error of the predicted effect (R)). Hashed bars reflect the experimentally measured effect of each combination of mutations tested. (C) The effect of recreating the most plausible evolutionary trajectory for 20_39 with (solid bars) or without (hashed bars) the addition of H215N. In all figures an un-paired t-test was used to determine whether there was a significant difference in chloramphenicol detoxification activity between two groups. (***, *p* ≤ 0.001; **, *p* ≤ 0.01; *, *p* ≤ 0.05). Data presented in all figures represent the average of at least four biological repeats ± 1 S.D.

Although it was the same residue, R225, that was substituted in each case, the substituting residues possessed very different chemical properties (negatively charged aspartate in 20_39 *versus* non-polar valine in 36_37). This, together with the observation that none of our top 30 selected variants had retained a basic residue at position 225 (Figure 1C), indicated that it was essential for arginine 225 to be eliminated before the other active site substitutions could make a discernible contribution to improved chloramphenicol detoxification.

We also found evidence of higher-order epistasis beyond the requirement for elimination of R225. For example, both evolved variants contained the substitutions S41Y and T219Y, neither of which conferred a significant improvement in chloramphenicol detoxification when introduced to NfsA individually (Figure 3A) or together (Figure 3B). When each was introduced into an R225V or R225D background, S41Y yielded a significant increase in chloramphenicol detoxification, but T219Y did not (Figure 3B). However, the combination of S41Y and T219Y together with R225V or R225D gave a further significant improvement (Figure 3B). Numerous examples of sign epistasis can also readily be observed in the full network diagram (e.g., the blue circles indicate a negative impact for certain combinations of substitutions; Figure 2D-E, Supplementary Figure S1). For example, H215N (present in variant 20_39) is detrimental to chloramphenicol detoxification activity when substituted into the R225D or R225D/S41Y backgrounds, and somewhat neutral in combination with R225D/S41Y/T291Y, but significantly enhances activity in combination with R225D/S41Y/T291Y/F227H (Figure 3C). Overall, our data suggest that complex epistatic interactions render >99% of the possible 5040 evolutionary pathways (that might be traversed from wild type NfsA to either 36_37 or 20_39) broadly inaccessible to iterative mutagenesis strategies.

### Improved chloramphenicol detoxification is underpinned by loss of activity with 1,4- benzoquinone

The hypothetical evolutionary trajectories depicted in Figure 2F highlight particularly pertinent intermediate combinations of mutations. We considered that interrogating the intermediate variants might shed light on the mechanistic basis of improved chloramphenicol detoxification. In particular, we wanted to determine how activity with a presumed native substrate like 1,4-benzoquinone was affected during the hypothetical evolutionary progression towards improved chloramphenicol detoxification. For this, the enzyme intermediates identified in the most probable stepwise evolutionary trajectory (Figure 2F) were purified as His-tagged proteins and *in vitro* kinetics assays were conducted with both chloramphenicol and 1,4-benzoquinone (Supplementary Table S1). From this data, it was evident that the first substitution of both hypothetical trajectories (i.e., the elimination of R225; Fig. 3A) was sufficient to abolish nearly all 1,4-benzoquinone activity, which was then unmeasurably low across all subsequent substitutions (Figures 4A and 4E). The sustained loss of 1,4-benzoquinone activity throughout each trajectory strongly suggests that this activity cannot co-exist with chloramphenicol reduction. This is reinforced by examination of the chloramphenicol detoxification activities of the complete set of hypothetical evolutionary intermediates (Supplementary Figure S1); the variants that retained R225 were on average no better than wild type NfsA at defending host cells against chloramphenicol (mean fold-improvement of 1.0 ± 0.5), while cells expressing variants that contained the substitution R225V or R225D were on average able to tolerate 4.0 ± 2.4 fold higher chloramphenicol concentrations than those expressing wild type *nfsA* (Table S2).

**Figure 4:**
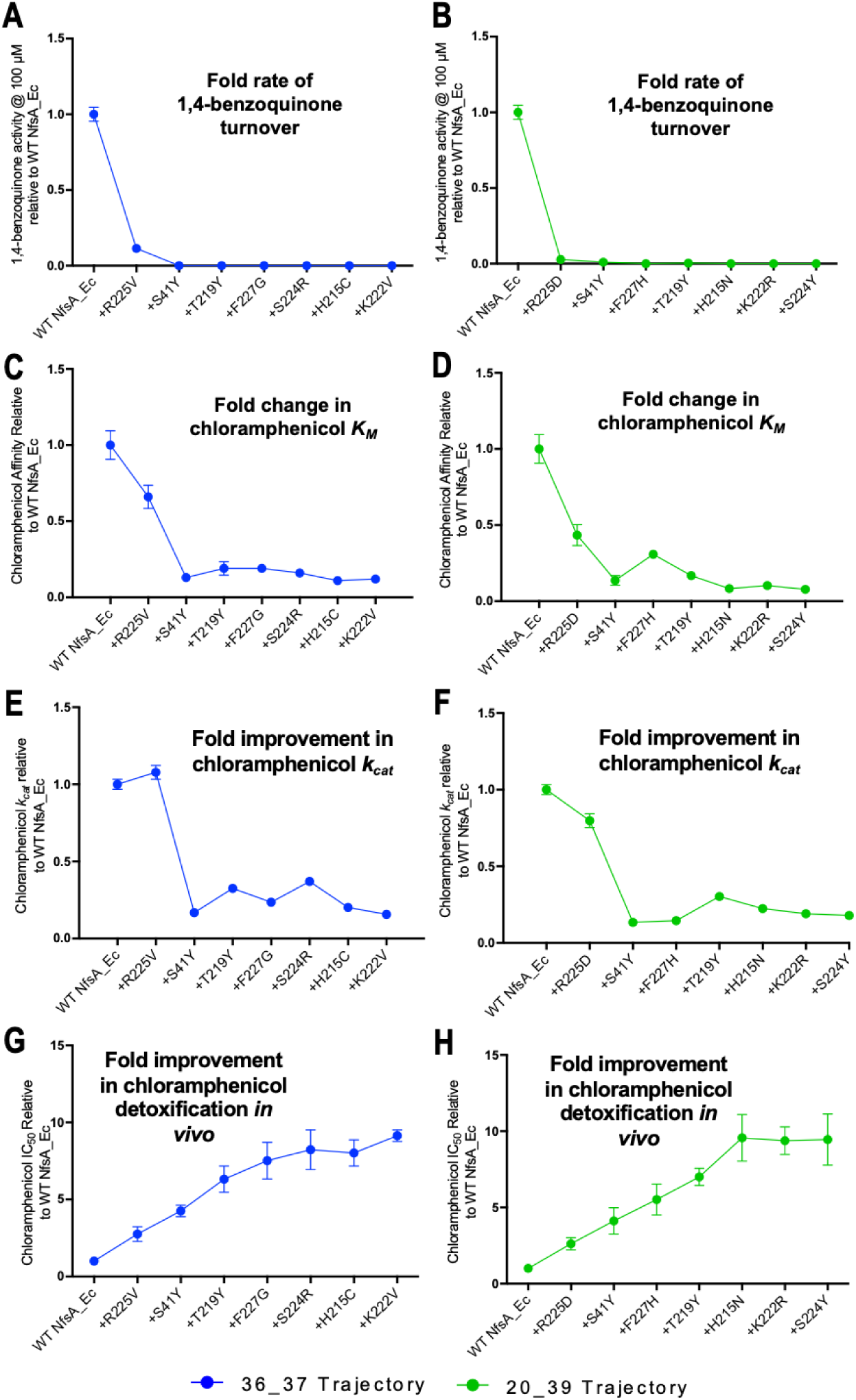
Activity analysis with 1,4-benzoquinone (A, B) and chloramphenicol (C-H) during the evolutionary progression of 36_37 (left, blue) and 20_39 (right, green). **(A, B)** Fold rate of turnover of 1,4- benzoquinone (starting concentration 100 µM, with 250 µM NADPH co-substrate) relative to wild type NfsA for each intermediate variant in the evolutionary trajectory of 36_37 (**A**) and 20_39 (**B**). (**C, D)** Fold change in chloramphenicol *K*_*M*_ relative to wild-type NfsA for each intermediate of 36_37 (**C**) and 20_39 (**D**). (**E, F)** Fold increase in chloramphenicol *k*_*cat*_ relative to wild-type NfsA for each intermediate of 36_37 (**E**) and 20_39 (**F**), reproduced for convenience from Figure 2F. (**G, H)** Fold improvement in chloramphenicol detoxification (EC_50_) conferred to *E. coli* 7NT host cells by each variant relative to wild-type NfsA. Full Michaelis-Menten kinetic parameters are shown in Supplementary Table S1. All *in vitro* data presented is the average of three technical repeats ± 1 S.D. and all *in vivo* data presented in the average of at least four biological repeats ± 1 S.D.

The substitution S41Y that came next in both trajectories yielded a profound improvement in chloramphenicol *K*_*M*_, but also diminished *k*_*cat*_ substantially (Figures 4B&C and 4F&G). We observed the same S41Y NfsA substitution in our previous PR-104A study, and concluded that this most likely enables planar stabilisation and stacking of nitroaromatic substrates between the isoalloxazine rings of flavin mononucleotide (FMN) and the introduced tyrosine (Copp *et al.*, 2017). It is likely that a similar phenomenon explains the improved affinity for chloramphenicol observed here, with the decrease in catalytic turnover also arising as a consequence of enhanced stabilisation of the Michaelis complex. The subsequent substitutions in each trajectory then act to ‘tune’ the system, exerting only minor effects on *k*_*cat*_, but overall yielding incremental improvements in chloramphenicol *K*_*M*_ that largely mirror the improved chloramphenicol detoxification observed *in vivo* (Figures 4C&E and 4D&F). SDS-PAGE analysis confirmed that the expression levels were consistent for each intermediate variant throughout the evolutionary progression, eliminating this as a variable exerting substantial influence on the relative activity levels *in vivo* (Supplementary Figure S2).

### Impact of evolving improved affinity for chloramphenicol on alternate substrates

Collectively, the above data suggest that selection of lead variants for enhanced chloramphenicol detoxification within an *E. coli* cellular context resulted in small gains in catalytic efficiency, driven by substantially improved affinity for chloramphenicol as a substrate, as well as a profound loss of activity for an endogenous competing quinone substrate. We were interested to discover the impact that this may have had on unselected promiscuous activities of NfsA. We therefore used EC_50_ growth assays to assess the sensitivities of *E. coli* 7NT strains individually expressing either NfsA, variants 36_37 or 20_39, or the hypothetical evolutionary intermediates thereof, to five structurally diverse nitroaromatic prodrugs (Figure 5). We anticipated that the loss of competitive inhibition by endogenous quinones might have generically enhanced activity with each of these prodrugs, resulting in heightened host cell toxicity. However, for four of the five prodrugs, host sensitivity was largely unchanged (EC_50_ within a range of 0.8 to 2-fold that of the NfsA- expressing strain) when expressing any of the variants (Figure 5C-F). The exception was metronidazole, for which all variants exhibited similar gains in activity to chloramphenicol, despite the two compounds sharing little structural similarity (Figure 5A-B). Moreover, the introduction of R225V or R225D substitutions into NfsA (which largely eliminate 1,4- benzoquinone activity; Figure 4A-B) did not improve reduction of all prodrugs, but only significantly enhanced activity with metronidazole and CB1954 (Student’s t-test; Figure 5B,D). We therefore concluded that our selection for enhanced chloramphenicol detoxification was not driven exclusively by loss of the competing quinone activity, as this would have tended to also enhance activity with other alternate substrates.

**Figure 5:**
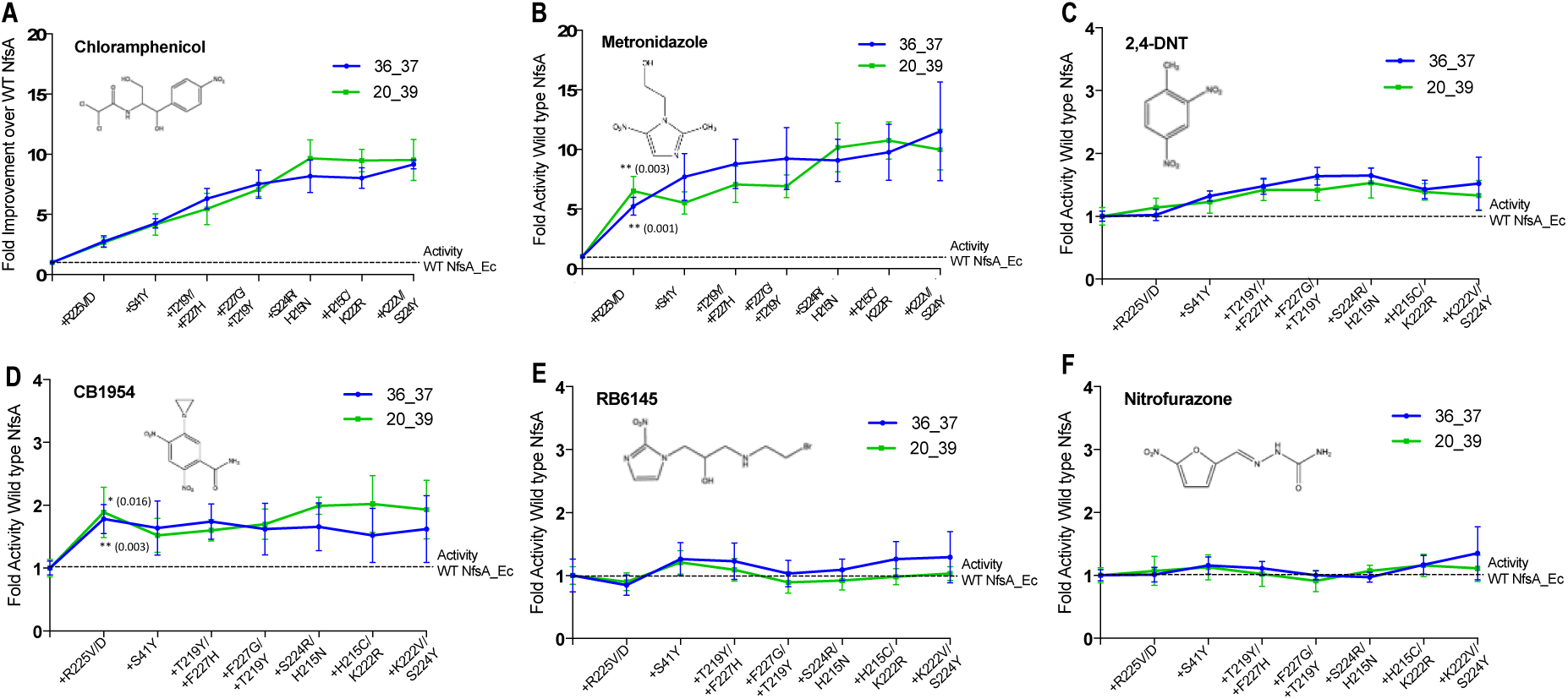
Activity analysis with nitroaromatic prodrugs during the evolutionary progression of 36_37 (blue) and 20_39 (green). *E. coli* 7NT cells expressing each of the hypothetical intermediate variants of 36_37 and 20_39 were tested in EC_50_ growth assays for (**A**) resistance to chloramphenicol; and (**B-F**) sensitivity to metronidazole, 2,4-DNT, CB1954, RB6145 and nitrofurazone, respectively. Data is presented as the fold improvement relative wild-type NfsA, with the average of four biological repeats ± 1 S.D. Chloramphenicol (**A**) and metronidazole (**B**) data are plotted on a different scale due to the large fold-improvements in activity. Where substitution of R225 caused a significant improvement in prodrug activation (Student’s t-test), the *p* value is noted in the figure panel (above the trendline for variant 20_39, and below for variant 36_37).

### Applications of 36_37 and 20_39 as dual selectable / counter-selectable marker genes

Whereas reduction of chloramphenicol is a detoxifying activity, reduction of metronidazole yields a toxic product. The serendipitous gains in metronidazole sensitivity that paralleled improved chloramphenicol detoxification inspired us to investigate whether these opposing activities might have useful molecular biology applications, by offering dual selectable and counter-selectable functionalities in a single gene. Counter-selectable markers, such as the *sacB* levansucrase gene from *Bacillus subtilis*, have multiple applications including the forced elimination of plasmids, and resolution of merodiploid constructs during allele exchange (Stibitz, 1994). However, they must typically be partnered with a selectable marker on the same DNA construct, to enable positive selection for the construct before its subsequent elimination. This occupies additional space, which is undesirable for size-restricted constructs, and means there is potential for the two genes to become separated by recombination events, leading to false positive or false negative outcomes.

Because metronidazole is cheap, widely-available, and has no measurable bystander effect in *E. coli* (*i.e.*, unlike many other nitroaromatic prodrugs its toxic metabolites are confined solely to the activating cell (Chan-Hyams *et al.*, 2018), we considered it ideally suited for counter-selection applications. We therefore tested the abilities of chloramphenicol to maintain, or metronidazole to force elimination of, plasmids bearing either 36_37 or 20_39 in *E. coli* 7NT. Cells were cultured for one hour in the absence of any selective compound, then plated on solid media amended with either 5 µM chloramphenicol or 10 µM metronidazole. The resulting colonies were then tested for retention or loss of the plasmid, respectively, with the expected outcome being realised in 100% of cases (94/94 colonies tested; Supplementary Figure S3). This suggests that our evolved variants might indeed have useful applications as dual selectable / counter-selectable marker genes.

## Discussion

By exploiting a powerful selection for antibiotic resistance, we were able to implement simultaneous mass-mutagenesis on an unprecedented scale, to amplify a promiscuous functionality. We acknowledge that this approach of focusing exclusively on eight key active-site residues means the reconstructed NfsA ‘engine’ is unlikely to be an optimal fit within the pre-existing chassis, and that further gains in activity would undoubtedly result by selecting residue substitutions in the second shell, or beyond. Nevertheless, we reasoned that our approach would allow us to gain comprehensive insight into key catalytic changes driving improved chloramphenicol detoxification, without being subject to the stochastic vagaries of error-prone PCR, or the well-established phenomenon that it can only access a limited and unbalanced repertoire of residues (on average, only 5.7 of the 19 alternative amino acids per codon position, with a bias toward similar residues (Hermes *et al.*, 1990). We also initially considered that this approach might allow us to leap to a fitness peak that iterative random mutagenesis strategies would be unable to scale. However, when we examined every hypothetical evolutionary intermediate, we discovered this was not the case; although stepwise evolution would have been greatly constrained in the progression from wild type NfsA to either of our top two variants, there were plausible trajectories to achieve these outcomes. Notably, the critical first step in any of these trajectories was the near-total elimination of the native quinone reductase activity, which was never restored and appears incompatible with the evolved activity. This is a very different scenario to the predominantly weak trade-offs observed in previous laboratory evolution studies (e.g., those reviewed by Kaltenbach *et al.*, (2016). Even the more recent work of Ben-David *et al.*, (2020) who encountered an abrupt activity trade-off when they evolved the calcium-dependent lactonase mammalian paraoxonase-1 into an efficient organophosphate hydrolase, found that the native functionality could subsequently be restored and was not incompatible with the evolved one (Ben-David *et al.*, 2013; Ben-David *et al.*, 2020).

By choosing to evolve an enzyme in its native cellular environment, we deliberately set out to explore the additional complexities of metabolic interference, which have potential to play dominant roles in shaping natural evolutionary outcomes. Copley recently described an equation that succinctly summarises how the rate of a promiscuous reaction in the presence of a native substrate might be improved by 1) increasing the concentration of the enzyme; 2) increasing the ratio of promiscuous to native substrate; and/or 3) altering the active site to diminish substrate competition, by enhancing binding or turnover of the promiscuous substrate, or decreasing binding of the native substrate (Copley, 2020). In a landmark 2015 study she and co-workers experimentally demonstrated the importance of diminished substrate competition, focusing on a single key Glu to Ala substitution that enabled several orthologs of ProA (L-gamma-glutamyl phosphate reductase, a key enzyme in proline synthesis) to replace *E. coli* ArgC (an N-acetyl glutamyl phosphate reductase required for arginine synthesis) (Khanal *et al.*, 2015). Where measurable, all of the substituted variants showed decreased affinity (increased *K*_*M*_) for the native substrate; and in all but one case there was substantial improvement in *k*_*cat*_*/K*_*M*_ for the promiscuous substrate as well (Khanal *et al.*, 2015). These findings are similar to our observation that elimination of quinone reductase activity from NfsA via substitution of R225 provided a platform for successive improvements in chloramphenicol affinity to amplify host cell resistance. Together, these examples support the proposal of Kaltenbach *et al.*, (2016) that during natural evolution of a promiscuous activity there is likely to be active selection against the original function, as well as our own supposition that most previous laboratory evolution studies have evaded this phenomenon by focusing on heterologous enzymes and/or exogenously applied substrates. Moreover, our observations that the unselected promiscuous activities of NfsA (reduction of a structurally diverse panel of prodrugs) were mostly unaffected is consistent with their central thesis, that positive selection alone does not lead to specialisation. The emerging picture is that evolution in the natural intracellular milieu involves both selection for the new function, and selection against the old.

An interesting difference between our scenario and that of the Copley team is that NfsA is far less essential to the fitness of its host cell than ProA (e.g., deletion of *nfsA* does not impair *E. coli* growth even under oxidative stress from heavy metal challenge (Ackerley *et al.*, 2004)). This means that when a new stress is encountered and a promiscuous function becomes essential, as we have modelled here, the enzyme can potentially evolve without necessitating gene duplication to preserve the original function. An apparent “freedom to operate” is manifest in the vast diversity of primary functionalities observed in the superfamily of nitroreductases that NfsA belongs to (which spans activities as divergent as quinone reduction, flavin reduction to power bioluminescence, flavin fragmentation, dehalogenation and dehydrogenation (Akiva *et al.*, 2017)). Although this contrasts with the prevailing Innovation–Amplification–Divergence (IAD) model for natural enzyme evolution (Bergthorsson *et al.*, 2007), it may not be an exceptional scenario – rather, as previously argued by Newton *et al.*, (2015) it is likely that only a minority of enzymes in a cell are under active selection pressure at any time, and redundancy in metabolic networks means that there is latent evolutionary potential that can be immediately tapped to adapt to stress without the requirement of rare and costly gene duplication events. That a single mutation may suffice to rapidly amplify a desirable promiscuous activity simply by eliminating native substrate competition confers substantial ‘robustness’ at a cellular level, even if it means that individual enzymes may not be as robust as previously considered.

## Methods

### Chemicals

Chloramphenicol, metronidazole, 2,4-dinitrotoluene and nitrofurazone were purchased from Sigma-Aldrich. CB1954 was purchased from MedKoo Biosciences. RB6145 was synthesised in-house at the Ferrier Institute, Victoria University of Wellington.

### Simultaneous Site-Directed Mutagenesis Library Construction and Selection

To randomise the eight targeted residues of NfsA_Ec (S41, L43, H215, T219, K222, S224, R225 and F227) we designed a degenerate gene construct with NDT codons (specifying Arg, Asn, Asp, Cys, Gly, His, Ile, Leu, Phe, Ser, Tyr and Val) at all positions other than 219 (NHT codon, encoding Ala, Asn, Asp, His, Ile, Leu, Phe, Pro, Ser, Thr, Tyr and Val) and 222 (VNG codon, encoding Ala, Gln, Glu, Gly, Leu, Lys, Met, Pro, Thr, Val and two Arg codons). Initially a synthetic gene library was ordered from Lab Genius pre-cloned into plasmid pUCX (Prosser *et al.*, 2013), however this only yielded 15% of the 252 million unique variants in our final collection. The remaining 85% were generated ourselves by ordering the same sequence as a gene fragment library from GenScript and ligating it into pUCX at the *NdeI* and *SalI* restriction sites. The combined libraries were used to transform *E. coli* 7NT, a derivative of strain W3110 bearing gene deletions of seven endogenous nitroreductases (*nfsA, nfsB, azoR, nemA, yieF, ycaK* and *mdaB*) and the *tolC* efflux pump (Copp *et al.*, 2014). Electrocompetent *E. coli* 7NT cells were generated as per Sambrook and Russell (2001), and the transformation efficiency was enhanced using a yeast tRNA protocol modified from Zhu and Dean, (1999). Library selection was conducted on selective solid media containing LB agar supplemented with 100 µg.mL^-1^ ampicillin and either 45 or 47.5 µM chloramphenicol. Appropriate dilutions of the pooled library stock were spread over plates and incubated at 37 °C for 40 hours. Dilutions of the library were also spread over non-selective solid media (LB agar supplemented with 100 µg.mL^-1^ ampicillin) to estimate the number of transformants included in each selection. Enzyme intermediates of NfsA_Ec 36_37 and 20_39 were ordered as synthetic gene fragments from Twist Biosciences and subsequently ligated into the *NdeI* and *SalI* restriction sites of the vectors pUCX (for EC_50_ analysis) or pET28(a)^+^ (for purification of His_6_-tagged proteins).

### Growth Assays

For growth inhibition assays, a 96-well microtitre plate with wells containing 200 µL LB medium supplemented with 0.2% glucose (w/v) and 100 µg.mL^-1^ ampicillin was inoculated with *E. coli* 7NT nitroreductase strains and incubated at 30 °C with shaking at 200 rpm for 16 hours. A 15 µL sample of overnight culture was used to inoculate 200 µL of induction media (LB supplemented with 100 µg.mL^-1^ ampicillin, 0.2% (w/v) glucose and 50 µM IPTG) in each well of a fresh microtitre plate, which was then incubated at 30 °C, 200 rpm for 2.5 hours. Aliquots of 30 µL apiece from these cultures were used to inoculate four wells of a 384-well plate (two wells containing 30 µL induction media and two wells containing 30 µL induction media supplemented with 2 × the desired chloramphenicol concentration). The cultures were incubated at 30 °C, 200 rpm for 4 hours. Cell turbidity was monitored by optical density at 600 nm prior to drug challenge and 4 hours post challenge. The percentage growth inhibition was determined by calculating the relative increase in OD_600_ for challenged versus control wells.

For EC_50_ growth assays, 100 µL of overnight cultures as above were used to inoculate 2 mL of induction media and incubated at 30 °C, 200 rpm for 2.5 hours. A 30 µL sample of each culture was added to wells of a 384-well plate containing 30 µL of induction media supplemented with 2 × the final prodrug concentration. Each culture was exposed to 7-15 drug concentrations representing a 1.5-fold dilution series of drug and one unchallenged (induction media only) control. The cultures were incubated at 30 °C, 200 rpm for 4 hours. Cell turbidity was monitored by optical density at 600 nm prior to drug challenge and 4 hours post challenge. The EC_50_ value of technical replicates was calculated using a dose-response inhibition four-parameter variable slope equation in GraphPad Prism 8.0. The EC_50_ values of biological replicates were averaged to provide a final EC_50_ value.

### Evolutionary Trajectory Analysis

Full details of how the evolutionary trajectory analysis was conducted are available at https://github.com/MarkCalcott/Analyse_epistatic_interactions/tree/master/Create_mutation_network.

### Protein Purification and Steady-State Kinetics

Recombinant nitroreductases were cloned into the His_6_-tagged expression vector pET28(a)^+^, expressed in BL21 and purified as His_6_-tagged proteins. Enzyme reactions were carried out in 60 µL reactions in 96-well plates with a 4.5 mm pathlength. All reactions were performed in 10 mM Tris HCl buffer pH 7.0, 250 µM NADPH, an appropriate dilution of chloramphenicol or 1,4-benzoquinone substrate dissolved in DMSO (0-4000 µM chloramphenicol and 100 µM 1,4-benzoquinone), made up to volume with ddH_2_O. Reactions were initiated with the added of 6 µL of enzyme (8 µM or an appropriate concentration) and the linear decrease in absorbance was monitored at 340 nm measuring the rate of NADPH depletion as an indirect measured of substrate reduction. As neither chloramphenicol nor 1,4-benzoquinone interfere with the absorbance at 340 nm, the extinction coefficient of NADPH at 340 nm was used (chloramphenicol = 12,400 M^-1^cm^-1^ and *p-*benzoquinone = 6,220 M^-1^cm^-1^, as two molecules of NADPH are required to reduce chloramphenicol to the hydroxylamine form, while only one is required for the reduction of *p-*benzoquinone to the quinol). Technical replicates were plotted using Graphpad Prism 8.0 software and non-linear regression analysis and Michaelis-Menten curve fitting was performed.

### SDS-PAGE analysis for key intermediate NfsA variants

*E. coli* 7NT pUCX::*nfsA* variant strains were used to inoculate 200 μL LB media supplemented with 0.2% (w/v) glucose and Amp. Cultures were incubated overnight at 30 °C, 200 rpm. The next day, 100 μL of the overnight culture was used to inoculate 2 mL of LB induction medium (LB supplemented with 0.2% (w/v) glucose, Amp and 50 μM IPTG). Day cultures were grown at 30 °C, 200 rpm for 6.5 hours, after which the cultures were pelleted by centrifugation at 2500 × *g* for 5 min. The supernatant was decanted and the cell pellets resuspended in ∼100 μL of LB medium, after which the OD_600_ of a 1:100 dilution was measured. Cell cultures were normalised by dilution with additional LB medium so that a 1 in 100 dilution would give an OD_600_ reading of 0.1. A 12 μL sample of each culture was mixed with 5 × SDS loading buffer, heated at 95 °C for 5 min and subjected to SDS-PAGE analysis on a 15% acrylamide gel.

### Evaluating the selection / counter-selection potential of evolved *nfsA* variants

A single colony of an *E. coli* 7NT cells expressing *nfsA_*Ec 36_37 or 20_39 was used to inoculate a 3 mL overnight culture of LB supplemented with 100 µg.mL^-1^ ampicillin. The next day, 100 µL of each overnight culture was used to inoculate 10 mL fresh LB medium in a 125 mL baffled conical flask. The culture was grown at 37 °C, 200 rpm for 1 hour then the OD_600_ of the flask was determined. An appropriate dilution of each culture was plated on agar plates containing either LB-only, or LB amended with 10 µM metronidazole or 5 µM chloramphenicol. At 10 µM metronidazole, cells expressing NfsA_Ec 36_37 or 20_39 could not grow but cells bearing no plasmid could, while the reverse scenario applied with 5 µM chloramphenicol. Plates were incubated at 37 °C for 16 hours (LB-only or LB + metronidazole) or 40 hours (LB + chloramphenicol). To confirm the presence/absence of the plasmid bearing 36_37 or 20_39, 47 colonies from each condition were streaked on LB agar plates supplemented with 100 µg.mL^-1^ ampicillin and incubated at 37 °C for 16 hours, with growth indicating presence of the plasmid and no growth indicating absence of the plasmid. The same 47 colonies, alongside a negative control were further tested in a PCR screen with *nfsA*_Ec forward and reverse specific primers (Prosser *et al.*, 2013). A band approximately 720 bp indicated presence of the plasmid, while no band indicated absence of the plasmid.

### Statistical Analysis

Unless otherwise stated, data are given as the mean ± standard deviation. The software programme GraphPad Prism 8.0 was used for all statistical analyses. Differences between measured EC_50_ values of enzyme variants were determined by an unpaired Student’s t-test. A p-value of ≤ 0.05 was considered statistically significant with *** = p-value ≤ 0.001, ** = p-value ≤ 0.01 and * = p-value ≤ 0.05.

## Acknowledgements

We thank Professor Dan Tawfik for insightful suggestions on shaping the research and Associate Professor Nobu Tokuriki for his comments on an early draft of the manuscript. This research was funded by The Royal Society of New Zealand Marsden Fund (contract 15- VUW-037 to DFA and WMP, including a PhD scholarship for KRH). MHR received additional support from a Victoria University of Wellington (VUW) Doctoral Scholarship, RFL a VUW Masters Scholarship, and MJC from a research grant awarded by the Cancer Society of New Zealand (grant 18.05 to MJC and DFA).

## Supplementary Figures

**Supplementary Figure S1:**
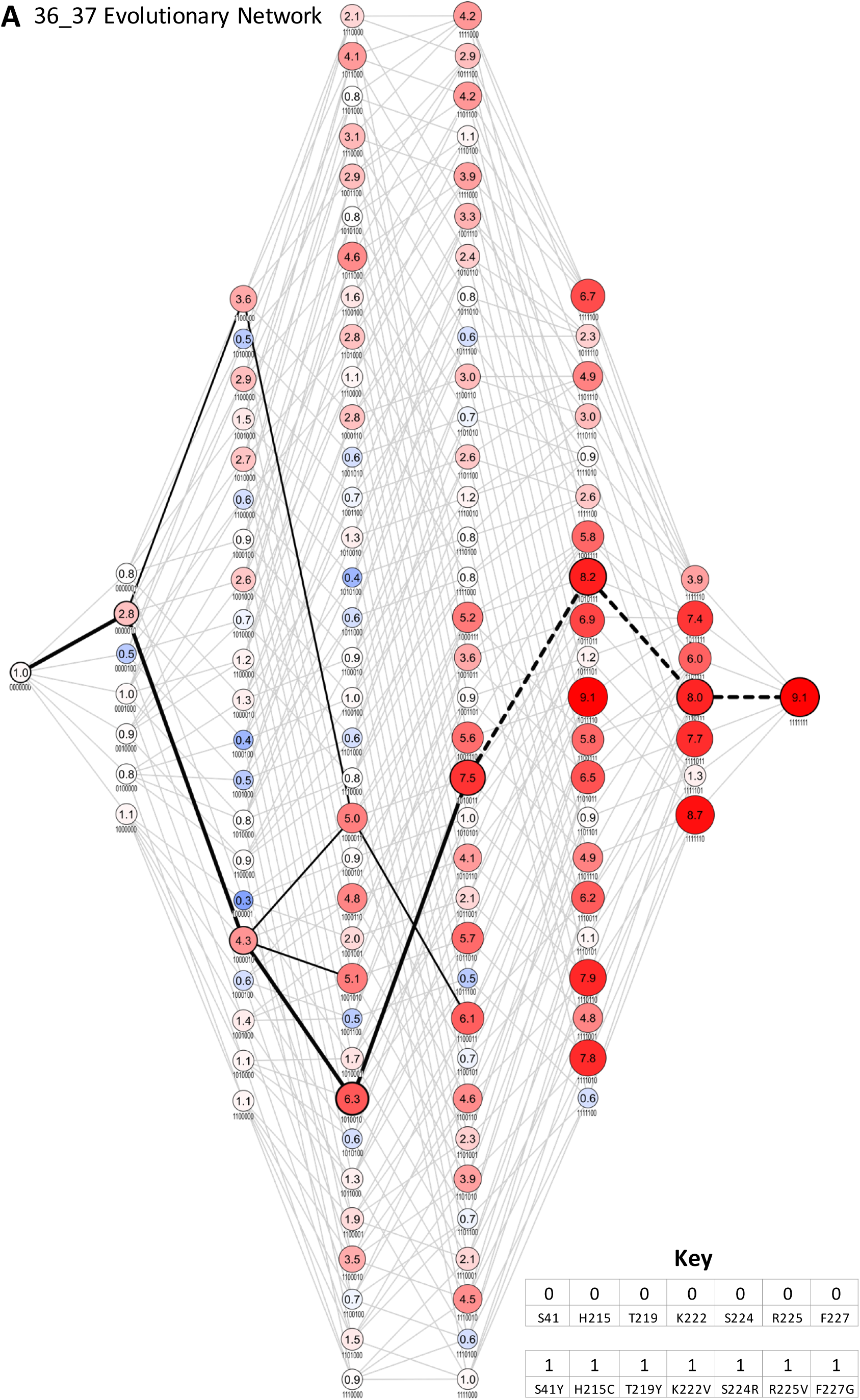
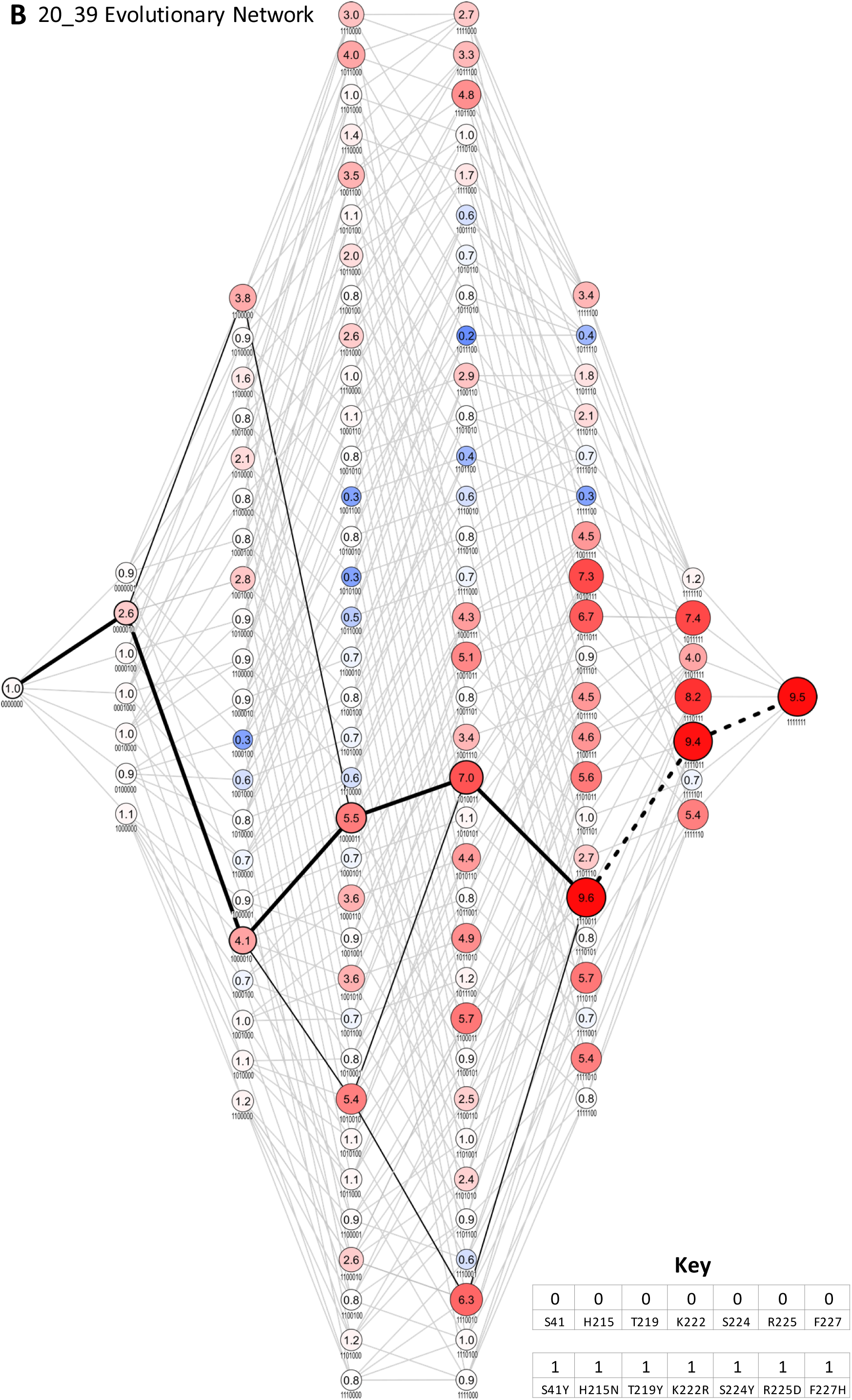
Evolutionary network of 36_37 and 20_39. Summary of all possible evolutionary trajectories for both (A) 36_37 and (B) 20_39. Each node represents an enzyme intermediate, with the substitutions present in the intermediate indicated by the presence (1) or absence (0) of the substitutions S41Y, H115C/N, T219Y, K222V/R, S224R/Y, R225V/D, F227G/H. respectively. The numbers within each node represent the fold activity relative to wild-type NfsA measured in *E. coli* EC_50_ assays; to facilitate rapid evaluation, the fold-activity for each node is also represented by the area of the circle, and by a colour-code (progressively less activity than wild-type NfsA in brighter shades of blue, and progressively more activity than wild-type NfsA in brighter shades of red). All 5040 possible evolutionary trajectories are marked by grey connecting lines. If a substitution resulted in a ≥ 16% increase in activity, the edge connecting the two nodes is shown in black (16% being greater than the average error for each set of 128 intermediates). The most probable stepwise mutagenesis pathway (shown in Figure 2D) is shown as thick black lines. A dotted line indicates that a substitution did not result in a ≥ 16% increase in activity, but formed part of the most probable stepwise mutagenesis evolutionary trajectory.

**Supplementary Figure S2:**
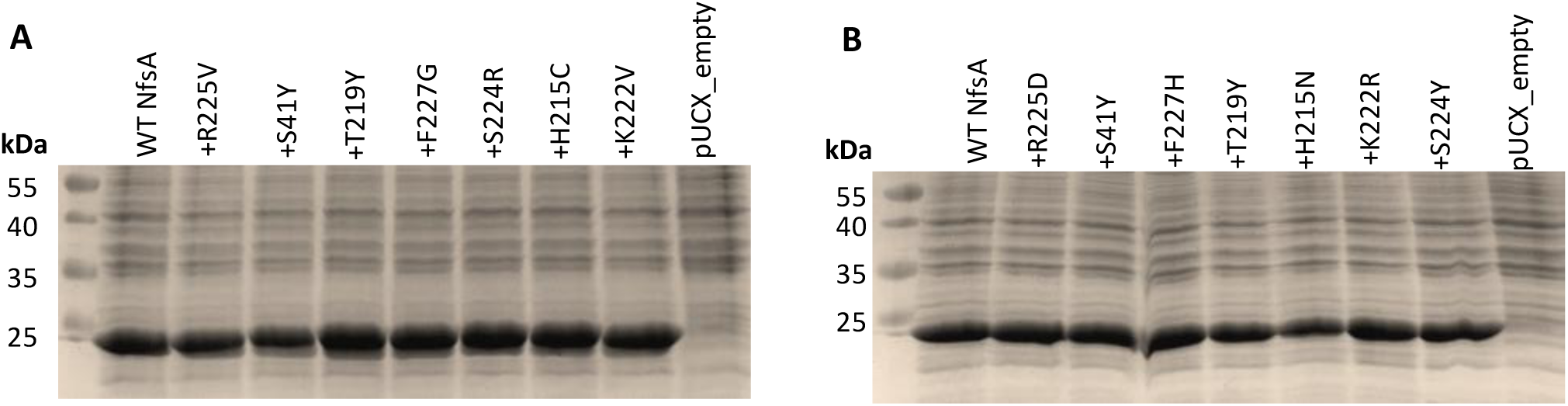
Relative enzyme expression levels for key intermediates from the most plausible trajectories for (A) 36_37 and (B) 20_39. Overnight *E. coli* 7NT cultures expressing pUCX::*nfsA* constructs were used to inoculate LB medium amended with 50 µM IPTG and incubated for 6.5 hours at 30 °C. Samples were normalised based on cell density (OD_600_). On each gel, lysate from the cells expressing wild-type *nfsA* was loaded in the left-most lane relative to the markers, then the strain expressing the R225V/D mutant, then the S41Y R225V/D double mutant, and so on, as indicated by the labels denoting each successive substitution. An *E. coli* strain bearing empty pUCX was included in the right-most lane as a negative control. The predicted molecular mass of NfsA is 26.8 kDa.

**Supplementary Figure S3:**
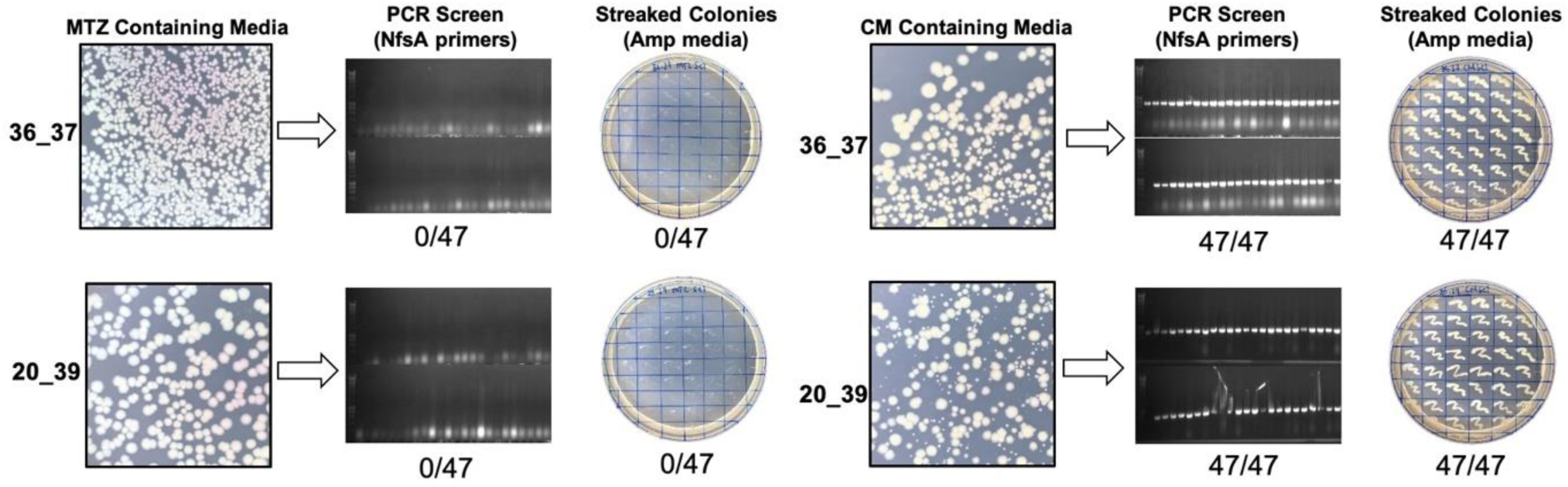
Growth of 36_37 and 20_39 on selective or counter-selective media. An overnight culture of 36_37 or 20_39 was added to a 1 in 100 dilution to a flask containing LB with no antibiotic or other selective compound. The culture was grown at 37 °C 200 rpm for 1 hour and then an appropriate dilution was plated over agar plates containing LB-only, 10 µM metronidazole (MTZ) or 5 µM chloramphenicol (CM). LB-only and MTZ plates were incubated at 37 °C for 16 hours, while CM plates were incubated for 40 hours. Forty-seven colonies from the MTZ and CM-containing plates, alongside a negative control were PCR screened with NfsA_Ec specific primers and streaked on media containing ampicillin (Amp) to provide a secondary indication of whether the plasmid bearing the nitroreductase gene was present.

**Supplementary Table S1:**
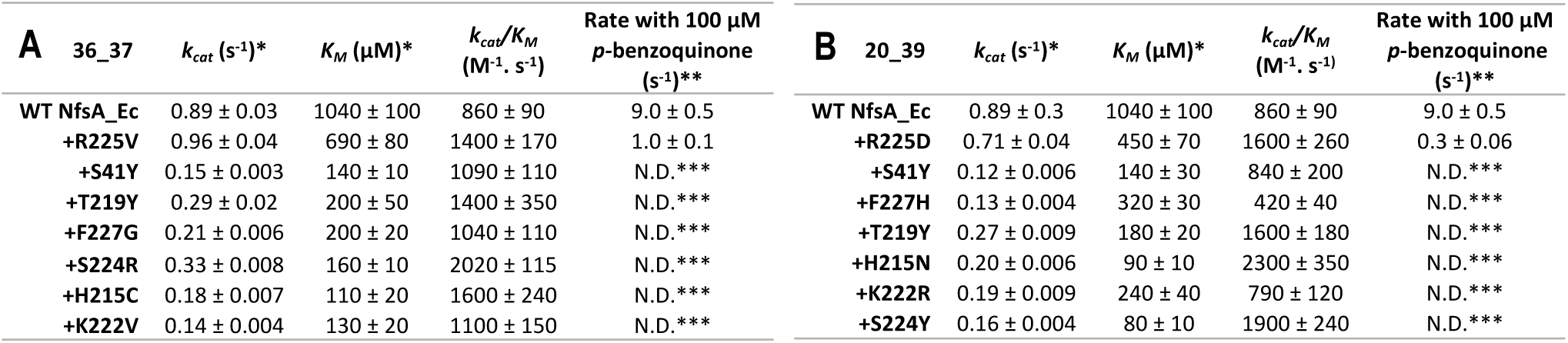
Kinetic parameters of chloramphenicol and 1,4-benzoquinone reduction for intermediates from the most plausible trajectories for (A) 36_37 and (B) 20_39. Apparent *K*_*M*_ and *k*_*cat*_ were calculated using Graphpad 8.0. Kinetic parameters could not be accurately determined for 1,4*-*benzoquinone, therefore the catalytic rate of 1,4-benzoquinone reduction was measured at a single high concentration of 1,4- benzoquinone (100 µM) with reactions initiated by addition of 250 µM NADPH. All reactions were measured in triplicate and errors are ± 1 S.D. In the left-most column, the terminology “+” refers to an enzyme variant that has the same amino acid sequence as the variant in the row above, plus the one additional substitution indicated. For example, “+R225V” describes a variant sharing an identical primary sequence to NfsA, with the additional substitution R225V. **\***Apparent *k*_*cat*_ and *K*_*M*_ as determined at 250 µM NADPH. **\*\***Measured rates following addition of 250 µM NADPH. **\*\*\***N.D. = not detectable (change in OD_340_ <0.1 s^-1^).

| A | 36_37 | Kinetic parameters | | | | Rate with 100 μM *p*-benzoquinone (s<sup>-1</sup>)** |
| --- | --- | --- | --- | --- | --- | --- |
| | | $k_{cat}$ (s<sup>-1</sup>)* | $K_M$ (μM)* | $k_{cat}/K_M$ (M<sup>-1</sup> s<sup>-1</sup>) | | |
| | WT NfsA_Ec | 0.89 ± 0.03 | 1040 ± 100 | 860 ± 90 | | 9.0 ± 0.5 |
| | +R225V | 0.96 ± 0.04 | 690 ± 80 | 1400 ± 170 | | 1.0 ± 0.1 |
| | +S41Y | 0.15 ± 0.003 | 140 ± 10 | 1090 ± 110 | | N.D.*** |
| | +T219Y | 0.29 ± 0.02 | 200 ± 50 | 1400 ± 350 | | N.D.*** |
| | +F227G | 0.21 ± 0.006 | 200 ± 20 | 1040 ± 110 | | N.D.*** |
| | +S224R | 0.33 ± 0.008 | 160 ± 10 | 2020 ± 115 | | N.D.*** |
| | +H215C | 0.18 ± 0.007 | 110 ± 20 | 1600 ± 240 | | N.D.*** |
| | +K222V | 0.14 ± 0.004 | 130 ± 20 | 1100 ± 150 | | N.D.*** |

**Supplementary Table S1: Kinetic parameters of chloramphenicol and 1,4-benzoquinone reduction for intermediates from the most plausible trajectories for (A) 36\_37 and (B) 20\_39.**
| B | 20_39 | Kinetic parameters | | | | Rate with 100 μM *p*-benzoquinone (s<sup>-1</sup>)** |
| --- | --- | --- | --- | --- | --- | --- |
| | | $k_{cat}$ (s<sup>-1</sup>)* | $K_M$ (μM)* | $k_{cat}/K_M$ (M<sup>-1</sup> s<sup>-1</sup>) | | |
| | WT NfsA_Ec | 0.89 ± 0.3 | 1040 ± 100 | 860 ± 90 | | 9.0 ± 0.5 |
| | +R225D | 0.71 ± 0.04 | 450 ± 70 | 1600 ± 260 | | 0.3 ± 0.06 |
| | +S41Y | 0.12 ± 0.006 | 140 ± 30 | 840 ± 200 | | N.D.*** |
| | +F227H | 0.13 ± 0.004 | 320 ± 30 | 420 ± 40 | | N.D.*** |
| | +T219Y | 0.27 ± 0.009 | 180 ± 20 | 1600 ± 180 | | N.D.*** |
| | +H215N | 0.20 ± 0.006 | 90 ± 10 | 2300 ± 350 | | N.D.*** |
| | +K222R | 0.19 ± 0.009 | 240 ± 40 | 790 ± 120 | | N.D.*** |
| | +S224Y | 0.16 ± 0.004 | 80 ± 10 | 1900 ± 240 | | N.D.*** |

**Supplementary Table S2:**
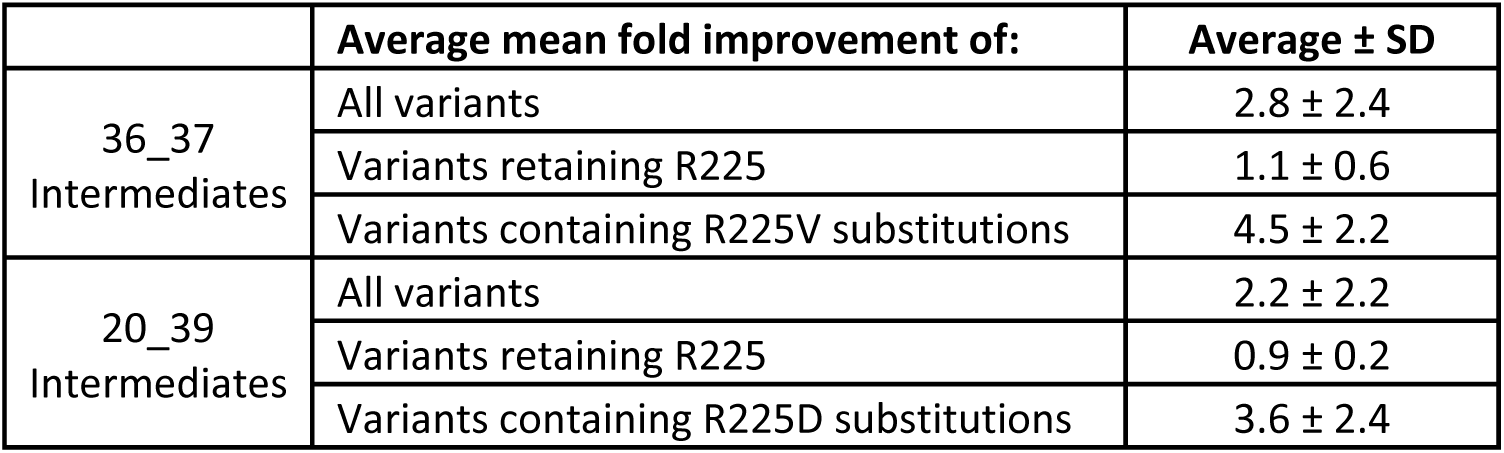
Average mean fold improvement for all NfsA_Ec variants that either retained R225 or contained a R225V/D substitution. To calculate the average fold improvement of variants retaining R225, the fold improvement relative to wild type NfsA_Ec of all 64 variants retaining R225 was averaged. To calculate the average fold improvement of variants with the R225V or R225D substitutions, the fold improvement relative to wild type NfsA_Ec off all 64 variants containing either R225V (in 36_37 intermediates) or R225D (in 20_39 intermediates) was averaged.

## References

Ackerley DF, Gonzalez CF, Keyhan M, Blake R 2^nd^, Matin A. 2004. Mechanism of chromate reduction by the *Escherichia coli* protein, NfsA, and the role of different chromate reductases in minimizing oxidative stress during chromate reduction. Environmental Microbiology 6:851–860. DOI: https://doi.org/10.1111/j.1462-2920.2004.00639.x

Aharoni A, Gaidukov L, Khersonsky O, Mc Gould S, Roodveldt C, Tawfik DS. 2005. The ‘evolvability’ of promiscuous protein functions. Nature Genetics 37:73–76. DOI: https://doi.org/10.1038/ng1482

Akiva E, Copp JN, Tokuriki N, Babbitt PC. 2017. Evolutionary and molecular foundations of multiple contemporary functions of the nitroreductase superfamily. Proceedings of the National Academy of Sciences USA 114:e9549–9558. DOI: https://doi.org/10.1073/pnas.1706849114

Ben-David M, Soskine M, Dubovetskyi A, Cherukuri KP, Dym O, Sussman JL, Liao Q, Szeler K, Kamerlin SCL, Tawfik DS. 2020. Enzyme Evolution: An Epistatic Ratchet versus a Smooth Reversible Transition. Molecular Biology Evolution 37:1133–1147. DOI: https://doi.org/10.1093/molbev/msz298

Ben-David M, Wieczorek G, Elias M, Silman I, Sussman JL, Tawfik DS. 2013. Catalytic metal ion rearrangements underline promiscuity and evolvability of a metalloenzyme. Journal of Moleulcar Biology 425:1028–1038. DOI: https://doi.org/10.1016/j.jmb.2013.01.009

Bergthorsson U, Andersson DI, Roth JR. 2007. Ohno’s dilemma: evolution of new genes under continuous selection. Proceedings of the National Academy of Sciences USA 104:17004–17009. DOI: https://doi.org/10.1073/pnas.0707158104

Chan-Hyams JVE, Copp JN, Smaill JB, Patterson AV, Ackerley DF. 2018. Evaluating the abilities of diverse nitroaromatic prodrug metabolites to exit a model Gram negative vector for bacterial-directed enzyme-prodrug therapy. Biochemical Pharmacology 158:192–200. DOI: https://doi.org/10.1016/j.bcp.2018.10.020

Copley SD. 2009. Evolution of efficient pathways for degradation of anthropogenic chemicals. Nature Chemical Biology 5:559–566. DOI: https://doi.org/10.1038/nchembio.197

Copley SD. 2015. An evolutionary biochemist’s perspective on promiscuity. Trends in Biochemical Sciences 40:72–78. DOI: https://doi.org/10.1016/j.tibs.2014.12.004

Copley SD. 2020. Evolution of new enzymes by gene duplication and divergence. The FEBS Journal 287:1262–1283. DOI: https://doi.org/10.1111/febs.15299

Copp JN, Mowday AM, Williams EM, Guise CP, Ashoorzadeh A, Sharrock AV, Flanagan JU, Smaill JB, Patterson AV, Ackerley DF. 2017. Engineering a multifunctional nitroreductase for improved activation of prodrugs and PET probes for cancer gene therapy. Cell Chemical Biology 24:391–403. DOI: https://doi.org/10.1016/j.chembiol.2017.02.005

Copp JN, Pletzer D, Brown AS, Van der Heijden J, Miton CM, Edgar RJ, Rich MH, Little RF, Williams EM, Hancock REW, Tokuriki N, Ackerley DF. 2020. Mechanistic understanding enables the rational design of salicylanilide combination therapies for Gram-negative infections. bioRxiv DOI: https://doi.org/10.1101/2020.04.23.058875

Copp JN, Williams EM, Rich MH, Patterson AV, Smaill JB, Ackerley DF. 2014. Toward a high-throughput screening platform for directed evolution of enzymes that activate genotoxic prodrugs. Protein Engineering, Design and Selection 27:399–403. DOI: https://doi.org/10.1093/protein/gzu025

Crofts TS, Sontha P, King AO, Wang B, Biddy BA, Zanolli N, Gaumnitz J, Dantas G. 2019. Discovery and characterization of a nitroreductase capable of conferring bacterial resistance to chloramphenicol. Cell Chemical Biology 26:559–570. DOI: https://doi.org/10.1016/j.chembiol.2019.01.007

Hall BG. 2004. Predicting the evolution of antibiotic resistance genes. Nature Reviews Microbiology 2:430–435. DOI: https://doi.org/10.1038/nrmicro888

Hermes JD, Blacklow SC, Knowles JR. 1990. Searching sequence space by definably random mutagenesis: improving the catalytic potency of an enzyme. Proceedings of the National Academy of Sciences USA 87:696–700. DOI: https://doi.org/10.1073/pnas.87.2.696

Kaltenbach M, Emond S, Hollfelder F, Tokuriki N. 2016. Functional Trade-Offs in Promiscuous Enzymes Cannot Be Explained by Intrinsic Mutational Robustness of the Native Activity. PLOS Genetics 12:e1006305. DOI: https://doi.org/10.1371/journal.pgen.1006305

Kaltenbach M, Jackson CJ, Campbell EC, Hollfelder F, Tokuriki N. 2015. Reverse evolution leads to genotypic incompatibility despite functional and active site convergence. Elife 4:e06492. https://doi.org/10.7554/eLife.06492.001

Kaltenbach M, Tokuriki N. 2014. Dynamics and Constraints of Enzyme Evolution. Journal of Experimental Zoology Part B-Molecular and Developmental Evolution 322:468–487. DOI: https://doi.org/10.1002/jez.b.22562

Khanal A, McLoughlin SY, Kershner JP, Copley SD. 2015. Differential Effects of a Mutation on the Normal and Promiscuous Activities of Orthologs: Implications for Natural and Directed Evolution. Molecular Biology and Evolution 32:100–108. DOI: https://doi.org/10.1093/molbev/msu271

Khersonsky O, Tawfik DS. 2010. Enzyme Promiscuity: A Mechanistic and Evolutionary Perspective. Annual Review of Biochemistry 79:471–505. DOI: https://doi.org/10.1146/annurev-biochem-030409-143718

Kobori T, Sasaki H, Lee WC, Zenno S, Saigo K, Murphy MEP, Tanokura M. 2001. Structure and site-directed mutagenesis of a flavoprotein from *Escherichia coli* that reduces nitrocompounds - Alteration of pyridine nucleotide binding by a single amino acid substitution. Journal of Biological Chemistry 276:2816–2823. DOI: https://doi.org/10.1074/jbc.M002617200

Liochev SI, Hausladen A, Fridovich I. 1999. Nitroreductase A is regulated as a member of the soxRS regulon of *Escherichia coli*. Proc Natl Acad Sci USA 96:3537–3539. DOI: http://doi.org/10.1073/pnas.96.7.3537

Newton MS, Arcus VL, Patrick WM. 2015. Rapid bursts and slow declines: on the possible evolutionary trajectories of enzymes. Journal of the Royal Society Interface 12:20150036. DOI: https://doi.org/10.1098/rsif.2015.0036

O’Brien PJ, Herschlag D. 1999. Catalytic promiscuity and the evolution of new enzymatic activities. Chemical Biology 6:R91–R105. DOI: https://doi.org/10.1016/S1074-5521(99)80033-7

Parry R, Nishino S, Spain J. 2011. Naturally-occurring nitro compounds. Natural Product Reports 28:152–167. DOI: https://doi.org/10.1039/c0np00024h

Paterson ES, Boucher SE, Lambert IB. 2002. Regulation of the *nfsA* Gene in *Escherichia coli* by SoxS. Journal of Bacteriology 184:51–58. DOI: https://doi.org/10.1128/jb.184.1.51-58.2002

Patrick WM, Firth AE, Blackburn JM. 2003. User-friendly algorithms for estimating completeness and diversity in randomized protein-encoding libraries. Protein Engineering 16:451–457. DOI: https://doi.org/10.1093/protein/gzg057

Poelwijk FJ, Tanase-Nicola S, Kiviet DJ, Tans SJ. 2011. Reciprocal sign epistasis is a necessary condition for multi-peaked fitness landscapes. Journal of Theoretical Biology 272:141–144. DOI: https://doi.org/10.1016/j.jtbi.2010.12.015

Prosser GA, Copp JN, Mowday AM, Guise CP, Syddall SP, Williams EM, Horvat CN, Swe PM, Ashoorzadeh A, Denny WA, Smaill JB, Patterson AV, Ackerley DF. 2013. Creation and screening of a multi-family bacterial oxidoreductase library to discover novel nitroreductases that efficiently activate the bioreductive prodrugs CB1954 and PR-104A. Biochemical Pharmacology 85:1091–1103. DOI: https://doi.org/10.1016/j.bcp.2013.01.029

Ramos JL, Gonzalez-Perez MM, Caballero A, van Dillewijn P. 2005. Bioremediation of polynitrated aromatic compounds: plants and microbes put up a fight. Current Opinion in Biotechnology 16:275–281. DOI: https://doi.org/10.1016/j.copbio.2005.03.010

Reetz MT, Kahakeaw D, Lohmer R. 2008. Addressing the numbers problem in directed evolution. ChemBioChem 9:1797–1804. DOI: https://doi.org/10.1002/cbic.200800298

Rich MH. 2017. Discovery and directed evolution of nitroreductase enzymes for activation of prodrufs and PET imaging compounds (PhD thesis). Victoria University of Wellington, Wellington, New Zealand.

Sambrook JF, Russell DW. 2001. Molecular cloning: a laboratory manual. Cold Spring Harbor Laboratory Press (eds). 3rd Edition. ISBN-100-87969-577-3

Smith AL, Erwin AL, Kline T, Unrath WCT, Nelson K, Weber A, Howald WN. 2007. Chloramphenicol is a substrate for a novel nitroreductase pathway in *Haemophilus influenzae*. Antimicrobial Agents and Chemotherapy 51:2820–2829. DOI: https://doi.org/10.1128/Aac.00087-07

Stibitz S. 1994. Use of Conditionally Counterselectable Suicide Vectors for Allelic Exchange. Methods in Enzymology 235:458–465. DOI: https://doi.org/10.1016/0076-6879(94)35161-9

Valiauga B, Williams EM, Ackerley DF, Cenas N. 2017. Reduction of quinones and nitroaromatic compounds by *Escherichia coli* nitroreductase A (NfsA): characterization of kinetics and substrate specificity. Archives of Biochemistry and Biophysics 614:14–22. DOI: https://doi.org/10.1016/j.abb.2016.12.005

Weinreich DM, Watson RA, Chao L. 2005. Perspective: sign epistasis and genetic constraint on evolutionary trajectories. Evolution 59:1165–1174. DOI: https://doi.org/10.1111/j.0014-3820.2005.tb01768.x

Williams EM. 2013. Development of bacterial nitroreductase enzymes for noninvasive imaging in cancer gene therapy (PhD thesis). Victoria University of Wellington, Wellington, New Zealand. DOI: http://hdl.handle.net/10063/4994

Williams EM, Little RF, Mowday AM, Rich MH, Chan-Hyams JV, Copp JN, Smaill JB, Patterson AV, Ackerley DF. 2015. Nitroreductase gene-directed enzyme prodrug therapy: insights and advances toward clinical utility. Biochemical Journal 471:131–153. DOI: https://doi.org/10.1042/BJ20150650

Winkler R, Hertweck C. 2007. Biosynthesis of nitro compounds. ChemBioChem 8:973–977. DOI: https://doi.org/10.1002/cbic.200700042

Yang G, Anderson DW, Baier F, Dohmen E, Hong N, Carr PD, Kamerlin SCL, Jackson CJ, Bornberg-Bauer E, Tokuriki N. 2019. Higher-order epistasis shapes the fitness landscape of a xenobiotic-degrading enzyme. Nature Chemical Biology 15:1120–1128. DOI: https://doi.org/10.1038/s41589-019-0386-3

Yunis AA. 1988. Chloramphenicol: relation of structure to activity and toxicity. Annual Review of Pharmacology and Toxicology 28:83–100. DOI: https://doi.org/10.1146/annurev.pa.28.040188.000503

Zhu H, Dean RA. 1999. A novel method for increasing the transformation efficiency of *Escherichia coli* application forbacterial artificial chromosome library construction. Nucleic Acids Research 27:910–911. DOI: https://doi.org/10.1093/nar/27.3.910

